# Unwrapping the ciliary coat: high-resolution structure and function of the ciliary glycocalyx

**DOI:** 10.1101/2024.09.28.615573

**Authors:** Lara M. Hoepfner, Adrian P. Nievergelt, Fabrizio Matrino, Martin Scholz, Helen E. Foster, Jonathan Rodenfels, Alexander von Appen, Michael Hippler, Gaia Pigino

## Abstract

The glycocalyx, a highly heterogeneous glycoprotein layer of cilia regulates adhesion and force transduction, and is involved in signalling. The high-resolution molecular architecture of this layer is currently not understood. We describe the structure of the ciliary coat in the green alga *Chlamydomonas reinhardtii* by cryo-electron tomography and proteomic approaches and present the high-resolution cryoEM structure of the main component, FMG1B. We describe FMG1B as a mucin orthologue which lacks the major *O*-glycosylation of mammalian mucins, but is *N*-glycosylated. FMG1A, a previously undescribed isoform of FMG1B is expressed in *C. reinhardtii*. By microflow-based adhesion assays we observe increased surface adhesion in the glycocalyx deficient double-mutant *fmg1a-fmg1b*. We find this mutant to be capable of surface-gliding, with neither isoform required for extracellular force transduction by intraflagellar transport. Our results find FMG1 to form a protective layer with adhesion-regulative instead of adhesion-conferring properties and an example of an undescribed class of mucins.

## Introduction

The green alga *Chlamydomonas* uses its cilia (also called eukaryotic flagella) as the primary means of motility and sensing of the external environment. In a liquid environment, *Chlamydomonas* cells swim by a breast-stroke motion. However, cells can also readily adhere to surfaces by their cilia, in which case the cilia are no longer beating. These cells are still able to move in a gliding fashion that is driven by intraflagellar transport (IFT). IFT is a system of polymeric trains moving along the microtubule based axoneme that constitutes the core of all cilia. The rapid motion of IFT trains in *Chlamydomonas reinhardtii* is sufficient to pull the surface-adhered cell along the cilia. While the exact binding of IFT trains to components of the ciliary membrane remains unclear, it has been shown that moving trains can move beads attached to the outside of cilia ^1^. As such, the outer coat of the cilia, generally referred to as the glycocalyx, must act as the main force transducing element between IFT and the surroundings. Consequently, the adhesion of the glycocalyx is of significant importance to gliding in *C. reinhardtii* and we have recently demonstrated that changes in *N*-glycosylation lead to reduced ciliary adhesion ^2^. In mammalian cilia, the glycocalyx of motile cilia is the main interactor with the mucosal layer covering the epithelial layer, but the precise nature of this interaction is currently unknown^3^.

Bloodgood and Sloboda have shown that the flagellar membrane glycoprotein 1B (FMG1B), an unusually large about 4545 amino-acid long molecule, constitutes a major part of the glycocalyx of *C. reinhardtii* ^4^, which localises to the ciliary coat and is visible as a densely stained layer in electron microscopy sections. However, the mutants used by Bloodgood have exhibited occasional instability of the phenotype with detectable levels of FMG1 atypical for *C. reinhardtii* insertional mutants. This reversal has primarily been attributed to the fact that the insertion in those specific mutants is in the 5’UTR of FMG1B which does not necessarily guarantee a complete null of the protein. In addition to FMG1B, *C. reinhardtii* has a homologous gene FMG1A with high primary sequence similarity to FMG1B that has been assumed to be not expressed. Recent proteomic studies have found evidence for the expression of FMG1A ^2,5^, but have not further studied the second isoform. Here, we use a combination of cryo-electron microscopy (cryoEM), cryo-electron tomography (cryoET), proteomic analysis as well as fluorescence microscopy and microfluidics on CRISPR/Cas mutants to demonstrate that: 1) Both FMG1 isoforms are expressed in *C. reinhardtii*, 2) are structurally highly similar and localise to the ciliary coat in the same manner; 3) the lack of both isoforms significantly increases ciliary adhesion and induces excessive microsphere binding; 4) that gliding motility is not impaired by lack of both isoforms. From our results, we conclude that the gliding mechanism in *C. reinhardtii* is likely not mediated by a single species but can be understood as a holistic effect of the entire membrane surrounding IFT trains. We suggest further that FMG1 exemplifies a new class of mucins and acts primarily as a protective layer with adhesion-regulating rather than adhesion-conferring properties.

## Results

### The isoforms FMG1A and FMG1B are expressed on the surface of *C. reinhardtii cilia*

With the recent extension of the CliP library^6^ we were able to obtain a mutant line with an insertion in the coding sequence of FMG1B (LMJ.RY0402.183636), specifically in the 5th exon (see Fig S1A) and we have verified the presence of the CIB1 insert by PCR (see Fig S1B). Analysis of whole CC4533 *fmg1b* cells by LC-MS/MS confirms the complete absence of FMG1B (see Fig S1C). Importantly, unique peptides are detected for FMG1A in both *fmg1b* cells as well as the background CC4533 strain, confirming that the second isoform is expressed on a population level. The *fmg1b* strain readily adheres to glass surfaces in gliding configuration and is indistinguishable from the background by phase-contrast microscopy (see Fig S1D).

**Figure S1:**
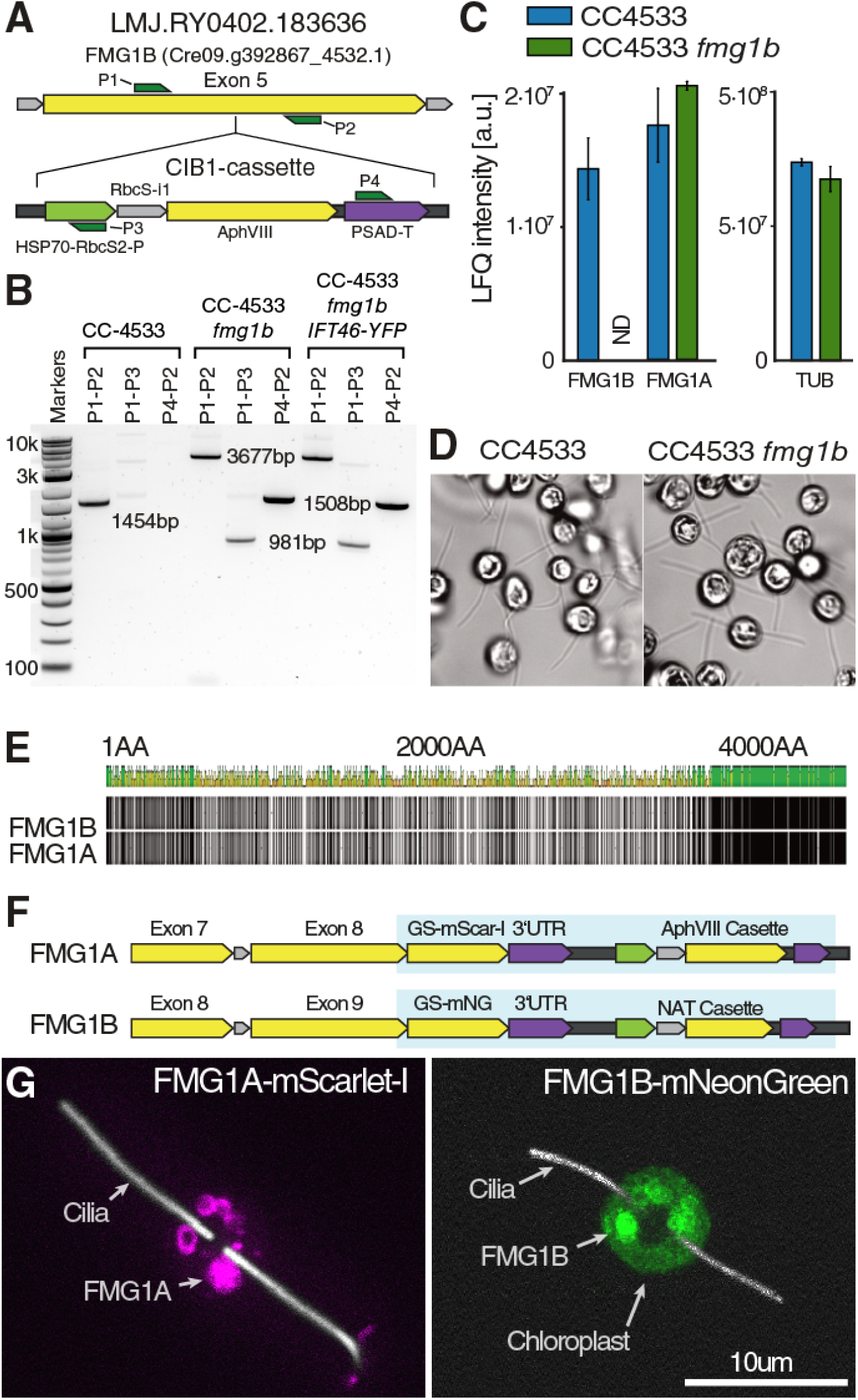
Detailed genetic and proteomic analysis of FMG1A and FMG1B. **A)** The CliP strain LMJ.RY0402.183636 (CC4533 *fmg1b*) has a CIB cassette insertion in exon 5 of FMG1B. **B)** PCR analysis confirms insertion of the CIB cassette by sizing and specific amplification. **C)** LC-MS/MS analysis confirms absence of FMG1-B in CC-4533 *fmg1b*, while protein intensity for FMG1A is slightly but not significantly increased. Relative abundance of βTubulin in wildtype and mutant is shown as control. **D)** Both CC4533 and CC4533 *fmg1b* adhere to glass surfaces in stretched cilia conformation, indicative of gliding motility. **E)** Amino-acid alignment shows significant primary sequence divergence of most of the n-terminal part of both FMG1 isoforms and a conserved c-terminal region. **F)** Genomic regions of CRISPR/Cas mediated endogenous fluorescent knock-ins for both FMG1 isoforms. Exogenous insert region is highlighted in blue. **G)** Spinning-disk confocal z-projections showing similar intracellular localization of both FMG1 isoforms to large, ring-like or vesicle-like structures. Cilia overlaid in intensity-rescaled white colour for reference. Scale bar is 10μm. Related to Figure 1.

In order to facilitate the observation of gliding in *fmg1b*, we crossed the cells to *ift46::NIT* IFT46-YFP cells and recorded the resulting cells in total internal reflection fluorescence (TIRF) microscopy. As shown in Fig. 1A, these cells readily glide in the absence of FMG1B. We then proceeded to investigate whether the lack of FMG1B causes a difference in ciliary adhesion strength. We assayed this property by subjecting adhered cells in a microchannel to a fluid flow while observing the surface of the microchannel by bright-field microscopy (see Fig. S2A). The drag force from the flow results in cells being detached over time (see Fig. S2B) and we measure the number of detached cells as a proxy for the adhesion strength. As such, we do not observe a significant difference in adhesion between CC4533 and the *fmg1b* mutant cells, indicating that the adhesion is not solely mediated by FMG1B. This led us to ask whether FMG1A could assume the same function as FMG1B, since both isoforms share an identical c-terminal tail region and have significant homology in the rest of the sequence (see Fig. S1F). To elucidate the 3D organisation and distribution of FMG1s in the glycocalyx, we performed cryoET on the intact cilia extending from frozen life *C. reinhardtii* cells. In wild-type cells, the ciliary membrane which surrounds the central axonemal structure is coated with a diffuse inner layer of low electron density and an outer, electron dense layer consisting entirely of FMG1 (see Fig. 1C-E). Importantly, in tomographic slices of CC-124 as well as CC4533 *fmg1b* cells, the same scale-like structure of the outer ciliary FMG1 coat can be observed (see Fig. 1D,E), confirming that both FMG1 isoforms assume the same ultrastructural arrangement in the glycocalyx. We then wondered to what extent either isoform is present in the cilia of wild-type cells. Since proteomic analysis can only measure populations and we were unable to distinguish the isoforms in cryoET, we generated the fluorescent dual knock-in line *FMG1A-mScarletI FMG1B-mNeonGreen* using CRISPR/Cas based precision genome editing (see Fig S1G). Observation of this line by spinning-disk confocal microscopy showed that either one of the two isoforms usually dominates, but the presence of both isoforms in the cilia can be occasionally observed in the ciliary membrane (see Fig. 1F). Further consolidating the similarity of the two FMG1 isoforms is a near identical localization of the newly produced protein inside the cell body (see Fig S1H).

**Figure 1:**
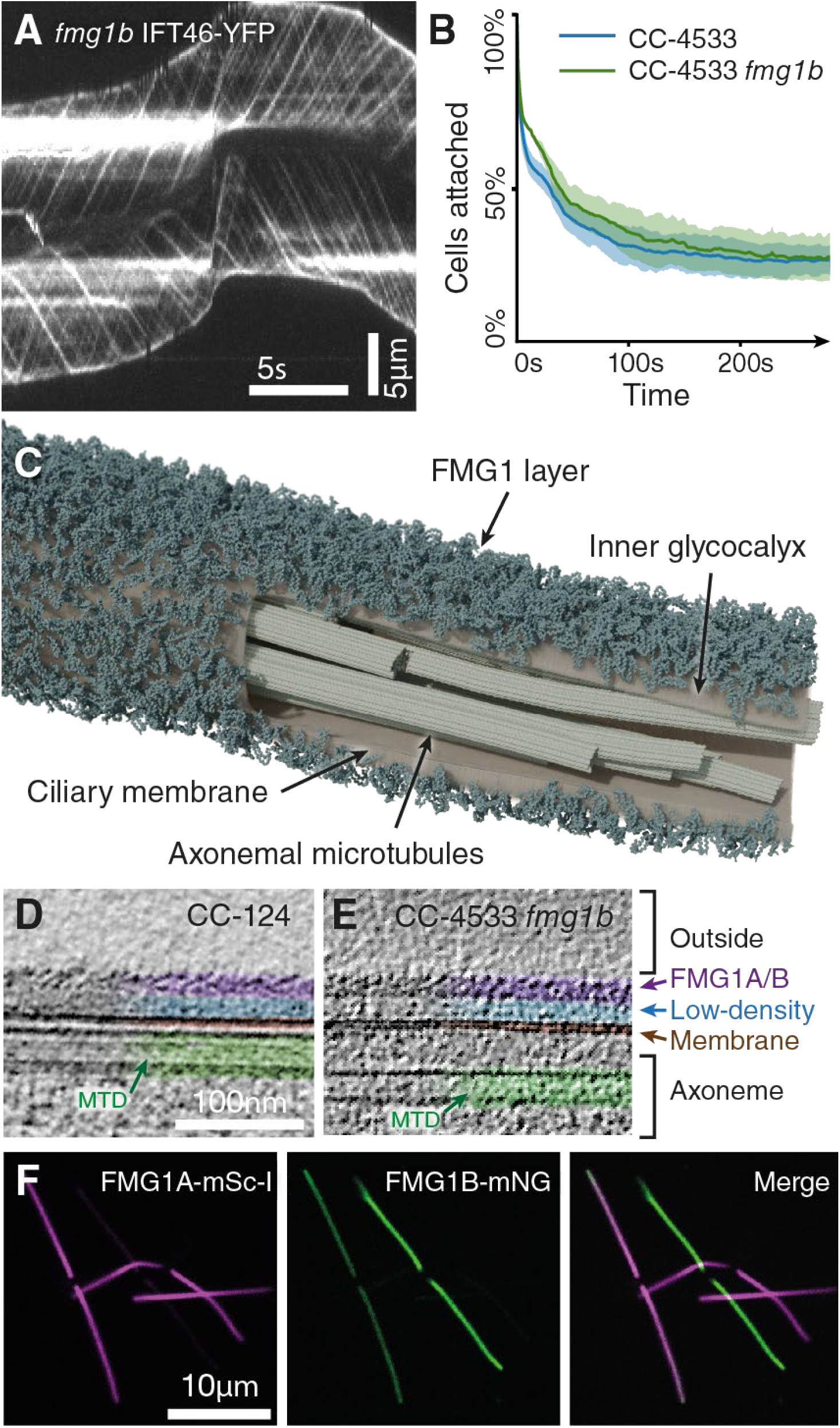
*C. reinhardtii* expresses both isoforms of the flagella membrane glycoprotein, FMG1-A and FMG1-B. **A)** Kymogram of the cilia of *fmg1b ift46*::NIT IFT46-YFP showing IFT mediated gliding in absence of FMG1B. Scale bars are 5s and 5μm. **B)** The absence of FMG1B does not cause loss of cellular adhesion to a polymer surface under 3.77ml/min flow. Average of three biological replicates. Shown error bars present standard deviation.**C)** Tomographic segmentation of a wild-type cilium with membrane partially hidden to show the ultrastructural organisation of the FMG1 layer coating the membrane which surrounds the central axonemal microtubules. **D)** Cryo-electron tomography (cryoET) slice of wild-type cilia showing the scale-like arrangement of FMG1 on the outer coat. Scale bar is 100 nm. **E)** CryoET slice of *fmg1* cilia with intact scale-like layer. **F)** Spinning disk confocal slices of adherent FMG1A-mScarlet-I FMG1B-mNeonGreen cells showing facultative co-expression of both FMG1 isoforms. Scale bar is 10μm. See also Figure S1.

**Figure S2:**
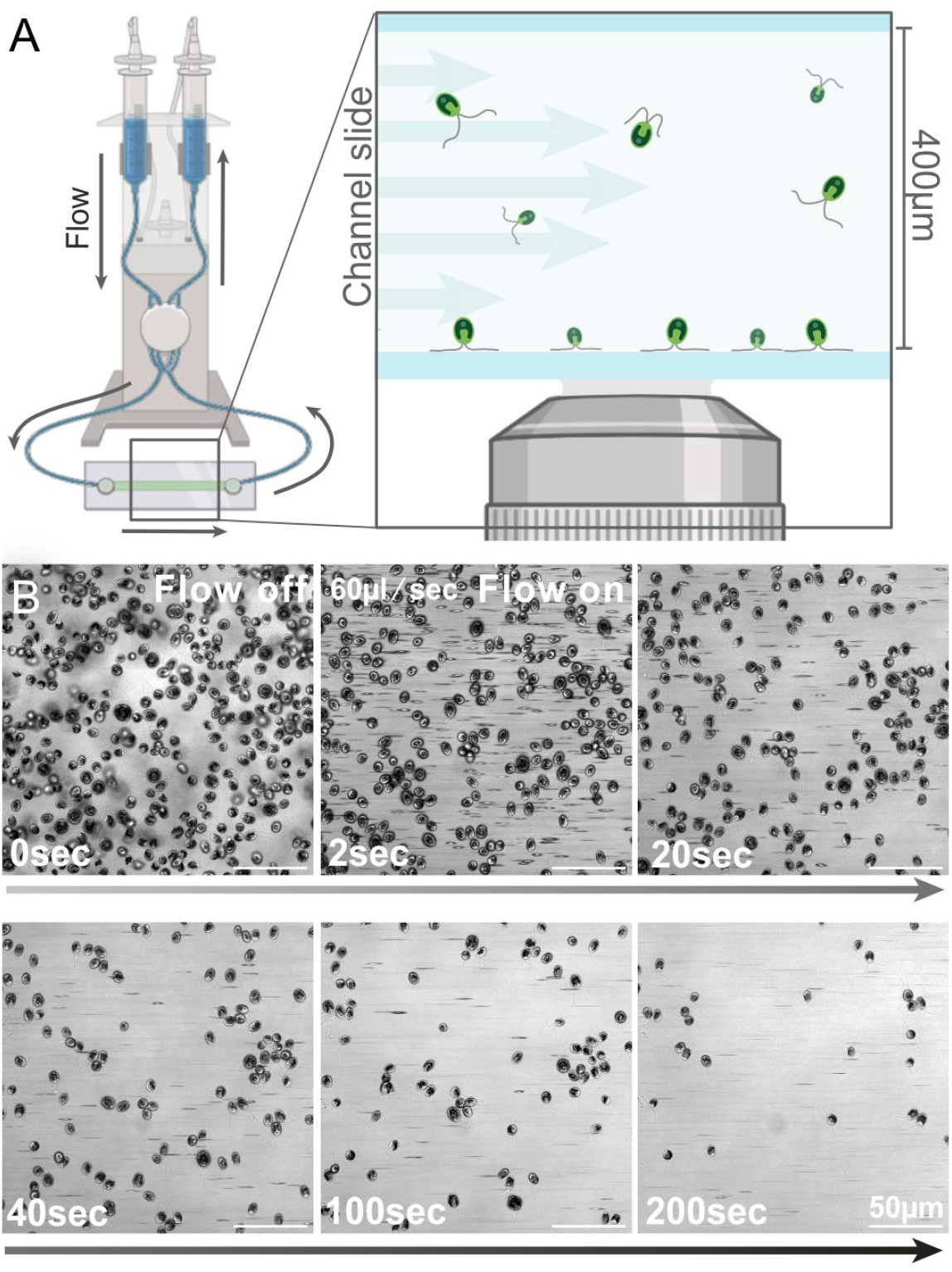
Microfluidic measurement for adhesion strength of attached *C. reinhardtii* cells to a microscopy slide. **A)** Schematic overview of the fluidic setup. Buffer is pumped from syringe reservoirs through a microscope slide with a microchannel by a peristaltic pump. **B)** Brightfield micrographs of adhered *C. reinhardtii* cells attached to the bottom polymer surface of the microchannel. Under flow, cells detach at a rate proportional to the strength of adhesion and can be seen moving as horizontal streaks through the image. Scale bar is 50μm. Related to Figure 1 and Figure 2.

### Depletion of both isoforms of FMG1 has no impact on gliding ability but increases cilia-substrate adhesion

Having established the presence of a second isoform, we generated a CRISPR/Cas mediated dual null mutant line CC124 *fmg1a-fmg1b* via an intermediate CC124 *fmg1b* line and set out to characterise this new line with regard to gliding motility and adhesion. Despite a total lack of FMG1, these cells adhered to a microscope slide with their cilia in gliding configuration (see Fig. 2A). To measure gliding activity, we crossed CC124 *fmg1a-fmg1b* to *ift46::NIT IFT46-mNeonGreen (IFT46-mNG)* to produce an IFT labelled but glycocalyx deficient line and subjected cells to gliding analysis using TIRF microscopy. Unexpectedly, these cells were still able to glide (Fig. 3B) with similar gliding velocities of 0.68µm/sec ± 0.78 in FMG1^+^ cells and 0.97µm/sec ± 1.00 in FMG1^−^ cells (Fig.2C). Likewise gliding distances were similar with a mean gliding distance of 5.2µm ± 2.4µm in FMG1^+^ cells vs 6.9µm ± 2.8µm in the FMG1^−^ line (Fig.2D). To compare the relative adhesion strength of CC124 and CC124 *fmg1a-fmg1b*, cells were reconstituted in calcium free Hepes/NMG buffer before transfer to the microchannel to encourage cells to settle on the surface. Under these conditions CC124 and CC124 *fmg1a-fmg1b* showed strong adhesion when exposed to stepwise increasing shear stress from 3.77ml/min to 51.94ml/min (Fig.2E). Surprisingly CC124 *fmg1a-fmg1b* cells showed a significantly slower detachment of cells, suggesting a significantly increased ciliary adhesion strength of glycocalyx deficient cells. To confirm this observation in an independent approach, we compared the capacity of the cilia of CC124 and CC124 *fmg1a-fmg1b* to bind microbeads: We mixed equal cell numbers of CC124 *fmg1a-fmg1b* and *FMG1A-mScarlet-I FMG1B-mNeonGreen* with polystyrene beads in one sample, thus ensuring the same conditions for both cell lines. While *FMG1A-mScarlet-I FMG1B-mNeonGreen* cilia were largely devoid of microspheres, excessive binding of beads could be observed along cilia of CC124 *fmg1a-fmg1b* accumulating primarily along the distal half of the cilium (Fig.2F). Considering these results, the absence of the major cilia glycocalyx seems to render the cilium more adhesive. As ciliary adhesion is sensitive to tunicamycin and altered *N*-glycosylation impacts adhesion force^2,7^, suggesting requirement of *N*-glycosylated proteins, we analysed protein abundance of confirmed ciliary *N*-glycoproteins^8^ that could compensate the loss of FMG1 in gliding and adhesion (Fig2G). Besides the uncharacterized membrane protein FAP170, no other known cilia *N*-glycoprotein showed significantly altered abundance.

**Figure 2:**
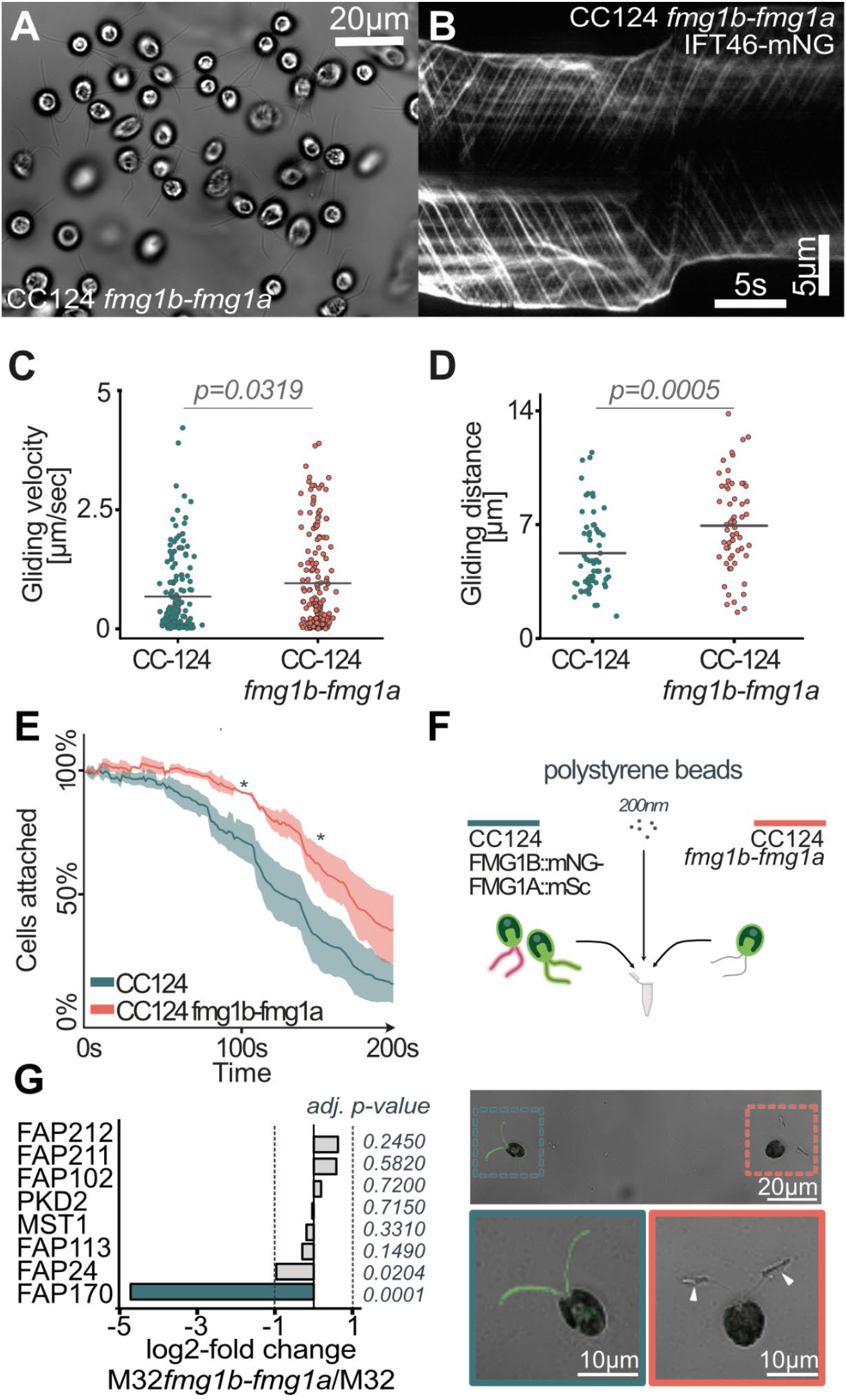
Depletion of FMG1 has no effect on gliding ability and renders cilia more adhesive. **A)** FMG1 deficient cells readily adhere in gliding configuration to glass coverslips. **B)** Kymogram of *fmg1b-fmg1a ift46::NIT IFT46-mNG* cilia showing IFT mediated gliding in absence of both FMG1 isoforms. Scale bars are 5s and 5μm. **C)** Detailed analysis of gliding velocities of surface adhered cells. (WT cells: 40; events:161, MT. cells: 42; events: 218) **D)** Gliding distance of 60 gliding events per strain analysed. **E)** Averaged detachment curves of CC124 and CC124 *fmg1a fmg1b* under stepwise increasing flow rate every 30 seconds (3.77ml/min-51.94ml/min). Three biological replicates were performed. Shown error bars present standard deviation. Two sample t-test was performed for statistical validation. **F)** Analysis of polystyrene microsphere binding capacity of CC124 *fmg1a fmg1b* and FMG1A-mScarlet-I FMG1B-mNeonGreen, shows excessive bead binding to cilia of CC124 *fmg1a fmg1b* (white arrow). G) Log2fold change(M32fmg1a-fmg1b / M32) of known *N*-glycosylated ciliary proteins quantified by LC-MS/MS.

**Figure 3:**
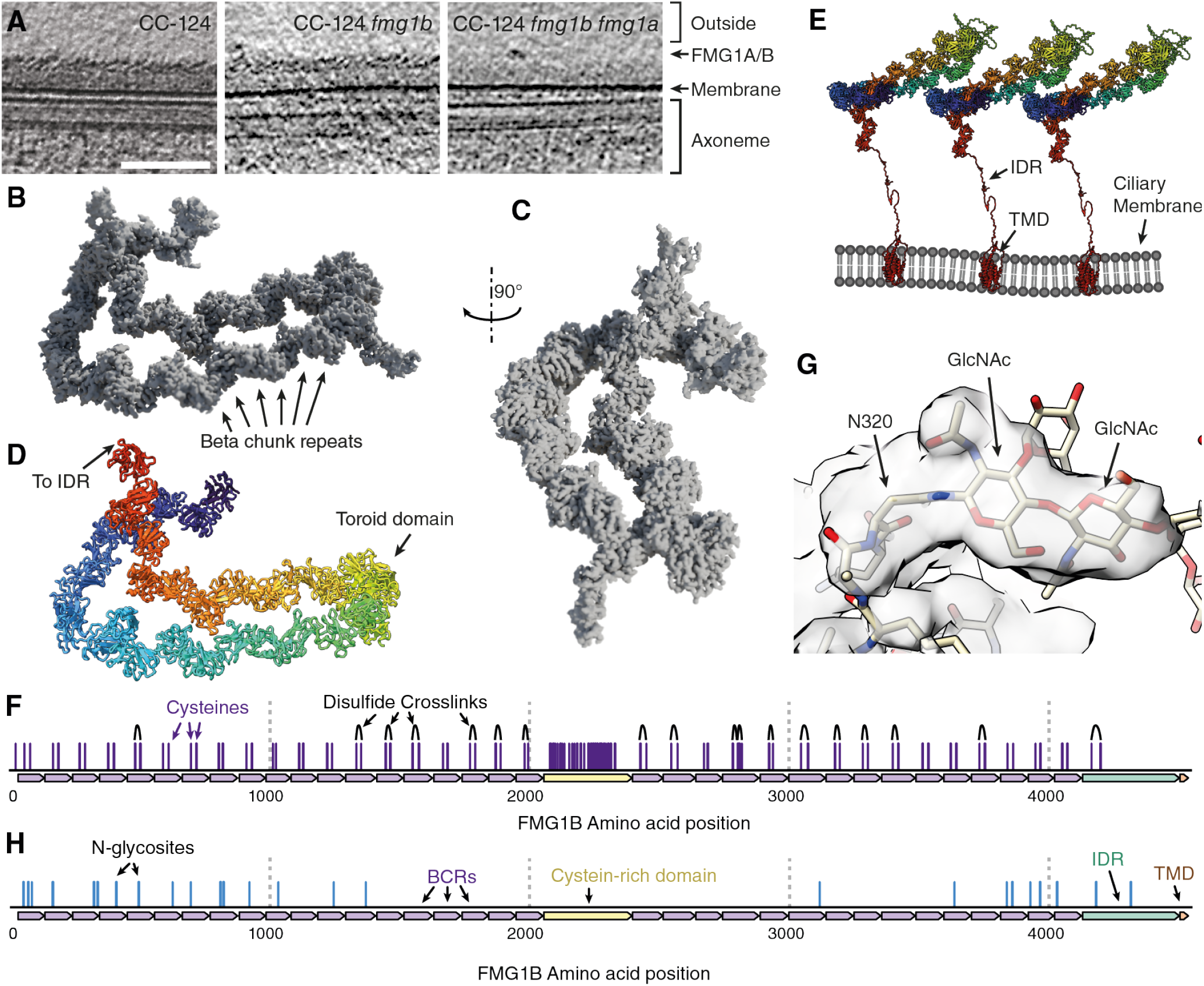
Cryo electron microscopy and structure of FMG1. **A)** CryoET slices showing the presence of the scale-like outer coat in cilia of WT CC124 cells as well as CC124 *fmg1b* cells. Cilia of double-ko CC124 *fmg1a fmg1b* cells lack the scale-like outer coat with only a diffuse inner coat layer present. **B)** Side view of cryoEM reconstruction of isolated FMG1B. Boxed area detailed in F). **C)** Front view of cryoEM reconstruction of isolated FMG1B. **D)** Integrative modellinging-based structure of FMG1B. **E)** Schematic representation of the organisation of FMG1 in the ciliary membrane. **F)** Map of FMG1B cysteine residues with disulfide bridges identified by structural proteomics. **G)** Representative detail view of Asn residue with bound N-glycan resulting in extra density. **H**) Map of N-glycosites identified by combined proteomics and cryo-EM analysis.

**Figure S3:**
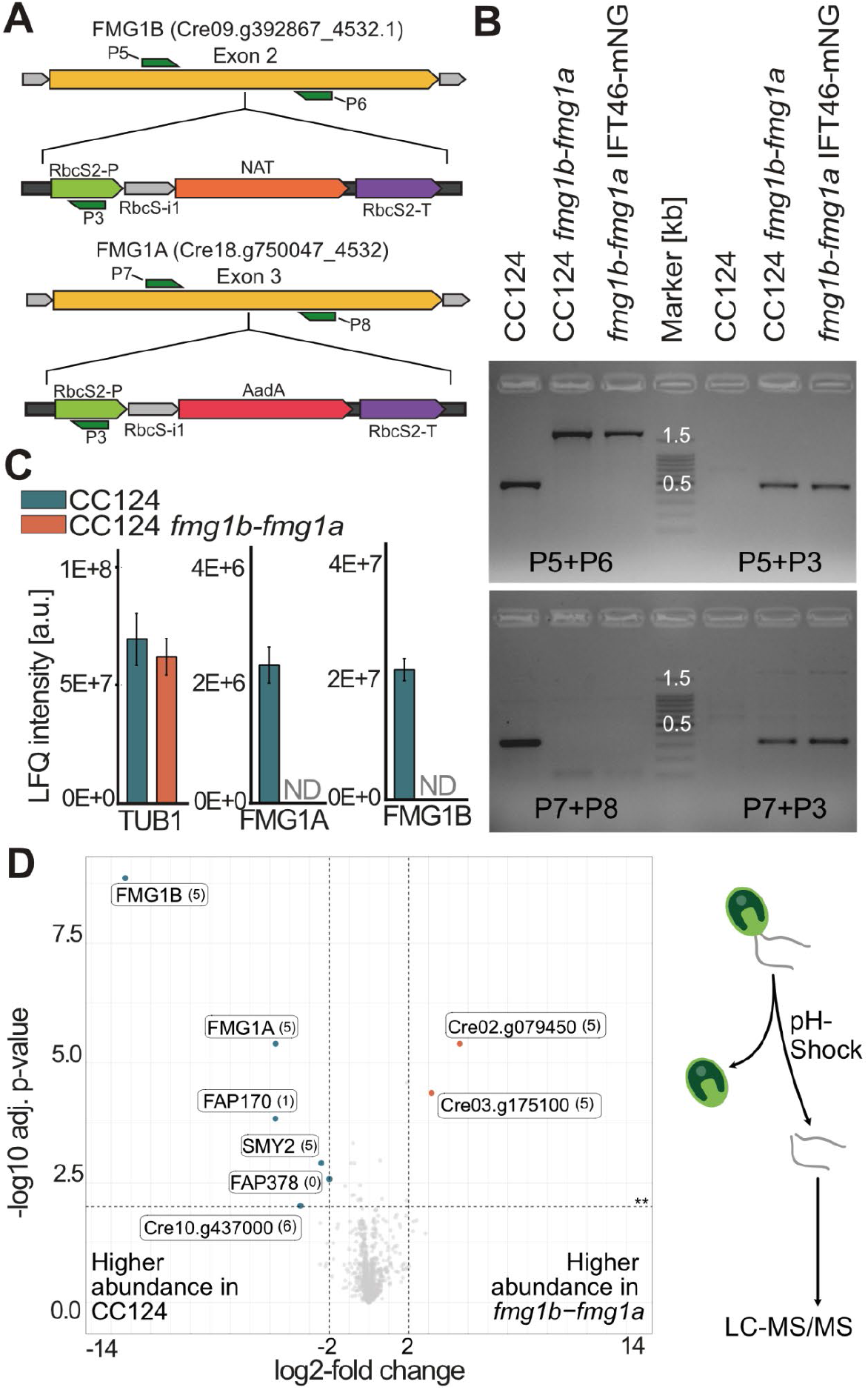
Genetic and proteomic analysis of CC124*fmg1a fmg1b*. **A)** Genomic maps of the glycocalyx deficient mutant, generated by CRISPR/Cas mediated insertion of NAT and AadA resistance cassettes in Exon2 and Exon3 of *fmg1b* and *fmg1a* respectively. **B)** PCR analysis confirming insertion of the cassette by sizing and specific amplification. **C)** LC-MS/MS analysis confirming the absence of FMG1-B and FMG1A in CC124 *fmg1a-fmg1b*. Relative abundance of βTubulin in wildtype and mutant is shown as control. **D)** Cilia proteome comparison of CC124 and CC124 *fmg1a-fmg1b* shows only minor changes in overall protein abundances. Bracketed numbers present replicates with imputed values (undetectable on peptide level). Data derives from five biological replicates. Related to Figure 2 and Figure 3.

### FMG1 is built from structural repeat with low sequence self-similarity

After establishing the phenotype of CC-124 *fmg1a-fmg1b* we then investigated the resulting ultrastructural changes to the glycocalyx. CryoET on the intact cilia of these lines revealed the absence of the scale-like layer that constitutes the outermost layer of the ciliary coat for the dual-knockout, but not for the intermediate CC124 *fmg1b* line (see Fig 3A). These data confirmed that the outer scale-like layer indeed consists of either isoform of FMG1 or a mix of both.

Having thus confirmed the identity of the scale-like outer coat, we set out to solve the high resolution structure of these unusual proteins. Due to the resolution limit of in-vivo cryoET we were not able to resolve a high-resolution structure of FMG1 directly from tomograms. Instead, we purified FMG1B from *C. reinhardtii* cilia by concanavalin A affinity chromatography and performed single particle cryoEM analysis.

The resulting ∼3Å structure of FMG1B revealed the molecule as a long, twisted chain of 34 individual globular subdomains of 109.6±10.9 amino acids and one cysteine rich coiled domain (Fig. 3B,C, Video S1). We generated a corresponding peptide model of FMG1B by an initial prediction with AlphaFold3^9^, followed by fitting with coarse-grained molecular dynamics^10^ and fine-tuning with ISOLDE^11^. The individual subdomains fold into a n Ig-like fold with each two orthogonal beta-sheets and one short alpha helix, which we called beta chunk repeats (BCR).

We modelled the semi-rigid structures composed of the chained BCRs to high confidence (see Fig. 3D), while the flexible nature of the cysteine rich domain resulted in a lower local resolution density which, together with the absence of bulky, recognizable amino acids, could not be assigned confidently. The c-terminus of FMG1 consists of an about 100 amino-acid long intrinsically disordered linker followed by a multi-helix transmembrane domain (TMD). Due to the inherent flexibility of the linker region, no densities could be resolved for either the linker region or the TMD. Using our integrative model of FMG1 we could explain the densities measured in cryoET by the rigid BCR structure making up the scales of the ciliary membrane which are anchored in the ciliary lipid bilayer by the TMD (see Fig 3E). Despite the high sequence variability between individual BCRs, every BCR contained a single highly conserved disulfide bridge as confirmed by MS assisted disulfide detection (Fig. 3F), while no disulfide bridges were detectable in the cysteine-rich domain under our conditions.Interestingly, we observe protruding densities by cryoEM that cannot be explained by the primary peptide sequence. Since FMG1 is known to be heavily glycosylated from biochemical analysis, we performed In-Source collision induced dissociation (IS-CID) MS/MS in combination with in-silico prediction^12^ to identify potential *N*-glycosites of FMG1B via mass spectrometry. Strikingly, we find near-perfect correlation between the identified candidate sites and the unexplained electron densities (see Fig. 3G and Table 1). As such, we were able to directly confirm 25 glycosylation sites of FMG1B, which were mostly located to the 16 n-terminal and the 9 c-terminal BCRs, all of which were in proximity to each other on one side of the FMG1B molecule (see Fig. 4H).

**Figure 4:**
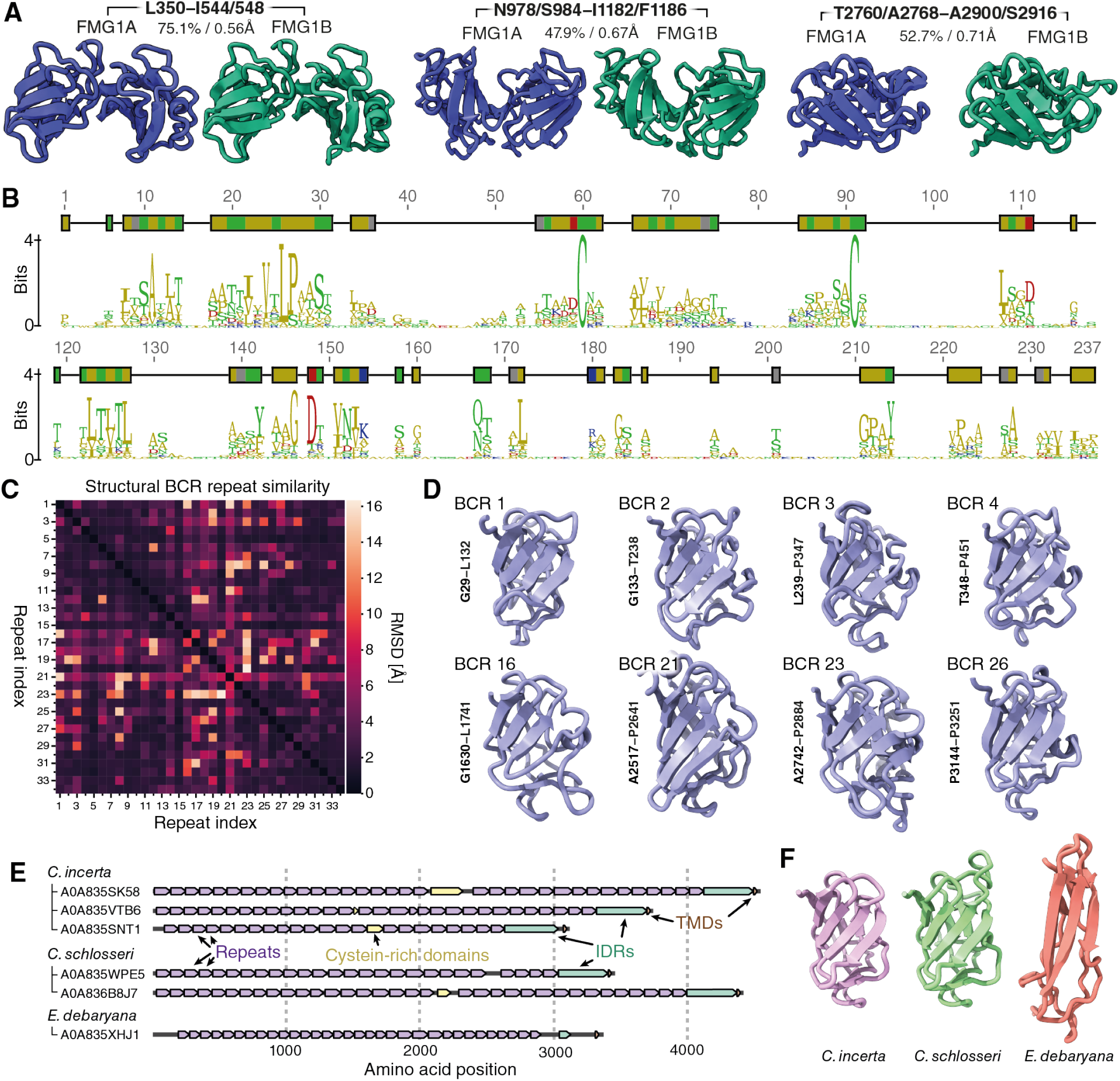
Structural characterization and conservation of FMG1 BCRs. **A)** Comparison of AlphaFold structures for FMG1A and FMG1B predict very high local structural homology despite significant sequence variability. Pairwise identity / RMSD values indicated for sequence and structural homology respectively. **B)** Sequence motif from all 34 aligned BCR repeats of FMG1B showing low overall sequence homology between BCRs except for a perfectly conserved pair of cysteines at positions 60 and 91. **C)** Pairwise root-mean square atomic displacement (RMSD) map for all 34 BCRs showing a high degree of structural homology between all BCRs in FMG1-B. **D)** Aligned example BCRs illustrating the conserved core beta-sheet fold and variable peripheral loops of different repeats. **E)** Structural annotation of transmembrane anchored homologues to FMG1 in three green algae *Chlamydomonas Incerta, Chlamydomonas Schlosseri* and *Edaphochlamys debaryana* showing examples of the wider family of the membrane-anchored folded-repeat based structure. **F)** Representative examples of the structural repeats annotated in **E)**.

We then used AlphaFold predictions as a tool to assess the structural similarity of the two FMG1 isoforms. Despite the fact that FMG1A and FMG1B only share about 61% pairwise identity, the protein fold of the isoforms was predicted to be almost identical, with typical root mean square deviation (RMSD) values of less than one angstrom between corresponding subdomains of either isoform (see Fig. 4A). These data suggested that FMG1A and FMG1B not only occupy the same space in the cilium, but also have almost the same protein structure. Likewise, the individual BCRs composing either isoform share little sequence homology (∼22% pairwise identity, Fig. 4B), but show a high degree of structural homology (Fig. 4C,D). Finally, suspecting that FMG1 might be just one example of a wider family of currently uncharacterized, related, proteins, we searched for homologues by linear alignment of individual domains and used AlphaFold predictions to verify potential hits. Unsurprisingly, the closely related *Chlamydomonas incerta* and *Chlamydomonas schlosseri*^13^ possess proteins that share high similarity, particularly in the fold of their repeat units (Fig. 4E,F). The number of repeats and the length—or presence—of a cysteine rich domain varied among these examples. Strikingly, despite a large sequence variance in the structural repeats, these proteins shared a much more conserved IDR domain. Based on this conservation, we repeated our search by focusing on the IDR domain and were able to find an example of a different repeat-fold in the more distant green alga *Edaphochlamys debaryana*^13^. These data suggest that FMG is indeed just one example of a highly variable family of mucin-like long and flexible proteins that are predicted to be targeted to the ciliary membrane of unicellular green algae.

## Discussion

Using cryoET, MS assisted protein quantification and a fluorescent dual knock-in approach we provide strong evidence that *C. reinhardtii* expresses FMG1-B and importantly FMG1-A. The glycoproteins localise to the cilium and to an unknown compartment within the cell body. Both isoforms, despite significant difference in amino acid sequence seem to obtain the same ultrastructure in vivo as revealed by the unchanged appearance of the ciliary coat in two independent FMG1-B knock-out mutants (Fig 1D,E, Fig 3A), while knock-out of both isoforms confirms that the outer ciliary glycocalyx is indeed solely composed of one or both of these two isoforms (Fig. 3A). While it is possible that other, unidentified components are localized to this layer and are lost in concert with the loss of FMG1, we do not detect additional unexplained densities in the outer coat at tomographic resolution. Loss of FMG1 is consistently correlated with a downregulation of FAP170 whose function is currently not known. Purifications of FMG1 from endogenous ciliary membranes do not contain detectable levels of FAP170 (in LC-MS/MS), indicating that there is no strong interaction between FAP170 and FMG1 and the partial loss of FAP170 in these mutants could be explained by translational regulation rather than a mechanically induced loss.

We suggest that FMG1 in *C. reinhardtii* is acting as a mucin orthologue. Mucins are ubiquitous in higher lifeforms and have evolved independently multiple times^14^. The protein family serves primarily in a protective and adhesion-regulatory function as a major component of the epithelial glycocalyx. Linear alignment of FMG1 to the human proteome matches the serine, proline and threonine rich regions to human Mucin 4. The diverse known mucins are classically characterised by a repetitive primary amino acid sequence and abundant carbohydrate modification. The repetitive primary sequence results naturally in a repetitive structural arrangement in known mucins. Conversely, both FMG1 isoforms consist of structural repeats of high similarity, but which share almost no primary sequence homology and as such raises the possibility of an entire class of currently not characterised mucin-like proteins that have been overlooked due to their lack of primary sequence repeats.

Interestingly, mucin 4 specifically has been shown to localise to motile cilia of the airways and anti-adhesive properties have been attributed to the protein due to an unspecific steric effect^15^. Both FMG1 and Muc4 contain a cysteine rich domain that is common to many mucins. The mechanochemical properties of known mucins arise due to the molecular interactions of long branched glycans^16^. While mammalian mucins are primarily decorated by *O*-glycosylation, the detectable glycans of FMG1 in *C. reinhardtii* are all *N*-linked. This isomorphism could be another indication that this new class of mucins in *C. reinhardtii* has evolved independently to other major lineages. Finally, while *O*-glycosylation is poorly characterised in *C. reinhardtii*, recent structural work on mastigonemes^17,18^ implies that only monosaccharide adducts are fused during *O*-glycosylation of serine and threonine which likely do not confer the same rheological properties as the long glycan chains typical of mucins.

Contrary to expectations, depletion of FMG1 increases the ability of cells to adhere to surfaces in our flow based and microsphere assays. This suggests that, in contrast to the current understanding of FMG1 as an adhesion promoter, the molecule likely acts as an adhesion regulator to avoid unintentional adhesion. Mutation of a gliding associated kinase GDAK has been implied to be sufficient to inhibit adhesion and microsphere binding in presence of FMG1^19,20^ and ciliary mediated adhesion is modulated by light, with blue-light inducing cells to adhere while red light induces detachment. Importantly, pronase treatment of cells under either light condition have further proposed protection of the peptidergic component required for adhesion under red-light^21^.

The ability to adhere to a surface is inherently required for gliding motility driven by IFT. Fundamentally, a single IFT train which is propelled along the axoneme has to transduce a force through the ciliary membrane and from there onto an extracellular surface to produce a relative motion. As the most abundant and the outermost positioned component of the ciliary coat, FMG1 has previously been thought to be a candidate to transduce this force^4^. We show here that the complete absence of the outer ciliary coat does not cause any significant change to the ability of a cell to move by gliding motility. This indicates that FMG1 is not required for force transduction through the membrane. We can not exclude that FMG1 together with other transmembrane components modulate IFT binding properties and act as force transducing elements.

The *N*-glycosylation inhibitor tunicamycin has been shown to impede cilia-microsphere interaction and adhesion forces are significantly reduced in mutants with altered *N*-glycan composition, suggesting that potential further adhesive candidates are likely to be *N*-glycosylated proteins^2,7^. Besides FMG1, eight ciliary membrane proteins have been shown to be *N*-glycosylated of which FAP113 and PKD2, likely in a complex with mastigonemes, have already been implicated in ciliary surface interaction^22,23^. Neither single knockouts of FAP113^22^ and PKD2/mastigoneme^23^ nor any combination of knock outs of FMG1A/FMG1B induced a detectable change in the ability of cells to glide it is therefore unlikely that either of these proteins is a key component of the gliding machinery. The remaining *N*-glycosylated ciliary proteins are namely: FAP170, FAP211, FAP212, FAP102, FAP24, and PKHD1^8^. Additionally, there likely exist further ciliary membrane proteins that we have not yet been able to identify with our current methods. Of particular interest among the identified candidates are FAP102, which contains two predicted fasciclin (FAS1) domains, and PKHD1. FAS1 domains are ancient cell adhesion molecules found in bacteria, yeast, plants and mammals^24^ and have been identified as putative cell-surface adhesion promoting proteins in the diatom *Phaeodactylum tricornutum*^25^. Finally, the transmembrane protein PKHD1 has potentially sufficient size to induce significant motions in the ciliary membrane/coat to induce gliding and additionally has a sizable predicted intraciliary domain of 541 amino acids which could potentially bind IFT and could therefore be an interesting candidate for future experiments.

In nature, *C. reinhardtii* cells likely navigate through water rich in particles such as sand, wood, and fibers^26^. Uncontrolled adhesion of such particles to cilia, as observed with polystyrene beads in this case, could significantly impair the motility of the cells. However, in environments like soil or silt, where water is confined to capillary microbridges between grains, the cells are likely capable of traversing dry patches by gliding when swimming is not an option. We conclude that FMG1 provides the essential adhesion-regulating properties needed to efficiently navigate through such diverse environments.

## Acknowledgments

The authors would like to acknowledge Dennis Diener and Samuel Lacey for their technical input and insights. We would also like to thank P. Suec from the Electron Microscopy Facility of Human Technopole. We would further like to acknowledge support from the MPI-CBG light microscopy facility as well as the MPI-CBG electron microscopy facility. A.P.N was supported by an EMBO Long–term fellowship under ALTF number 891-2018 as well as by an HFSP Cross-disciplinary fellowship with reference number LT000515/2019. We would like to acknowledge the European Research Council (ERC) under the European Union’s Horizon 2020 research and innovation program (grant agreement No. 819826). M.H. acknowledges funding from the Deutsche Forschungsgemeinschaft (DFG HI 739/12-4).

## Author contributions

Conceptualization, L.M.H, A.P.N, G.P, M.H; Methodology, L.M.H, A.P.N; Investigation, L.M.H, A.P.N, F.M, H.E.F; Data Curation, L.M.H, A.P.N; Visualisation, L.M.H, A.P.N; Formal Analysis, L.M.H, A.P.N; Writing – Original Draft,L.M.H, A.P.N; Writing – Review & Editing, L.M.H, A.P.N, G.P, M.H; Funding Acquisition, A.P.N, G.P, M.H; Resources, J.R, A.vA, G.P, M.H; Supervision, G.P, M.H

## Methods

Further information and requests for resources and reagents should be directed to and will be fulfilled by the corresponding authors.

## Materials Availability

- Strains and plasmids generated in this study have been deposited to the Chlamydomonas Resource Center. The collection numbers for the strains are listed in the KRT. These are also available from the lead author upon reasonable request.

## Data and code availability

- *All data reported in this paper will be shared by the lead contact upon request*. Original gel images of all main figures are provided as source data.
- The mass spectrometry proteomics deposited to the ProteomeXchange Consortium via the PRIDE (**Perez-Riverol et al**., **2019**) partner repository with the dataset identifier PXD055593. TIRF and confocal microscopy data used in this paper has been deposited to Zenodo with XXXXXX. FMG1-B peptide models have been deposited to RCSB PDB with the identifier 9GOS and single particle density maps have been deposited at EMDB with identifiers EMD-51499 (composite map), EMD-51661 (low resolution consensus map) and focused refinements: EMD-51649, EMD-51650, EMD-51654, EMD-51655, EMD-51657 and EMD-51658.
- Code used to analyse cryo tomography data is included in supplemental data file S1.
- Any additional information required to reanalyze the data reported in this paper is available from the lead contact upon request.

### Experimental Model and Study Participant Details

#### Chlamydomonas Cell Culture

Cells were grown photoheterotrophically in tris-phosphate-acetate (TAP) medium in dark:light cycle 8h:16h at 50 µmol photons*s^−1^*cm^−2^. Strains were maintained on 1.5% TAP agar plates and kept either in incubators under light or in aluminium coated trays under LED light bars.

### Method Details

#### CRISPR/Cas mutation

Mutants generated in this work were created by CRISPR/Cas assisted precision genome editing, following the methods described in our previous work^27,28^. Briefly, cells were grown on TAP 1.5% agar plates in 10h:14h dark:light under overhead LED illumination and harvested in active growth phase by scraping into liquid TAP medium. Cell walls were removed by three rounds of autolysin treatment, followed by three rounds of washing with TAP + 40mM sucrose. All centrifugation steps were performed at 700 RCF for 3 minutes. About 5 · 10^6^ cells were mixed with 5uL of 5μL of 5μM RNPs and 1.5μg of donor DNA, electroporated in an Invitrogen Neon Electroporator with three pulses of 2300V for 12ms, then dispensed into 1mL of TAPS in a 24 well plate for overnight recovery. Recovered reactions were plated on selective antibiotics (7.5μg/mL Nourseothricin, 100µg/mL Spectinomycin or 10µg/mL Paromomycin). Resulting colonies were picked into 96 well plates, grown up and prescreened with primers flanking the insertions by qPCR on a QuantStudio 7 Pro. Identified candidate insertions were then amplified by PCR and sequenced by Sanger or nanopore sequencing (Eurofins or Plasmidsaurus respectively). CC124-32M *fmg1b-fmg1a* was produced by an intermediate CC124-32M *fmg1b* step, while CC124-32M *FMG1A-mScarletI FMG1B-mNeonGreen* was produced in the background of CC124-32M *FMG1B-mNeonGreen* (CC-6012).

#### Cilia isolation

Cilia isolation from cultures in the mid-log growth phase was performed as described elsewhere by the pH shock method^29^. Pellets containing clean cilia samples were stored at −80°C until further use for sample preparation for mass spectrometric analysis.

#### Sample preparation for mass spectrometric analysis

Cilia and whole cell samples were reconstituted in lysis buffer (100mM Tris/HCl pH8, 2 % SDS, 1mM PMSF, 1mM Benzamidine) and subjected to sonication for 10 min. After pelleting cell debris, the protein concentration was determined using the bicinchoninic acid assay (BCA Protein Assay Kit by Thermo Scientific Pierce). Volumes corresponding to 60µg of protein were tryptically digested and desalted as described elsewhere^30^. For disulfide analysis cilia were reconstituted in lysis buffer+1% Igpal, reduction of proteins was omitted before digestion with trypsin or chymotrypsin.

#### Mass Spectrometry

Tryptic peptides were reconstituted in 2% (v/v) acetonitrile/0.1% (v/v) formic acid in ultrapure water and separated with an Ultimate 3000 RSLCnano System (Thermo Scientific). Subsequently, the sample was loaded on a trap column (C18 PepMap 100, 300 µm x 5 mm, 5 mm particle size, 100 Å pore size; Thermo Scientific) and desalted for 5 min using 0.05% (v/v) TFA/2% (v/v) acetonitrile in ultrapure water with a flow rate of 10 µL·min^−1^. Following, peptides were separated on a separation column (Acclaim PepMap100 C18, 75 mm i.D., 2 mm particle size, 100 Å pore size; Thermo Scientific) with a length of 50 cm. General mass spectrometric (MS) parameters are listed in table 3.

**Table 3:**
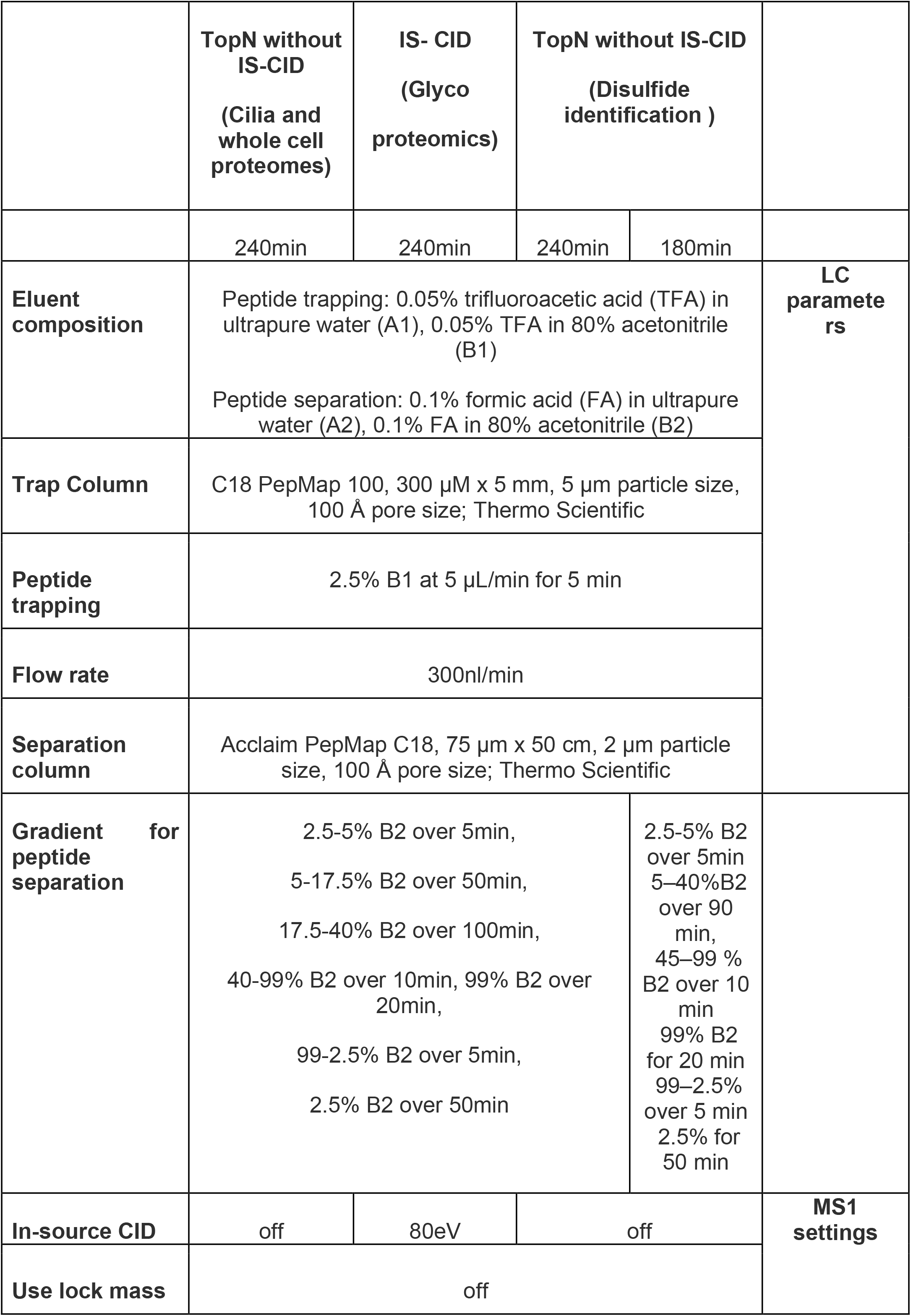
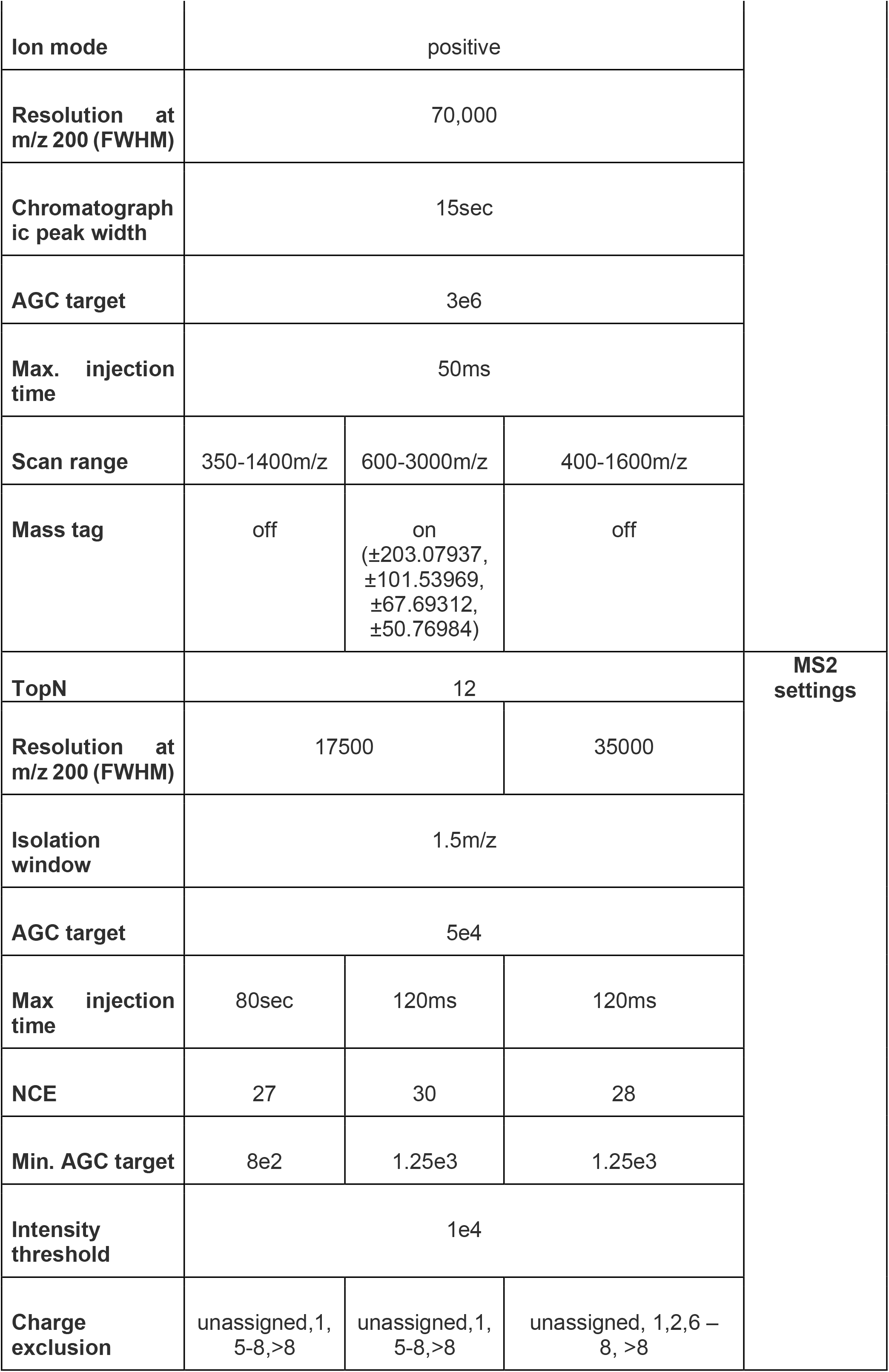
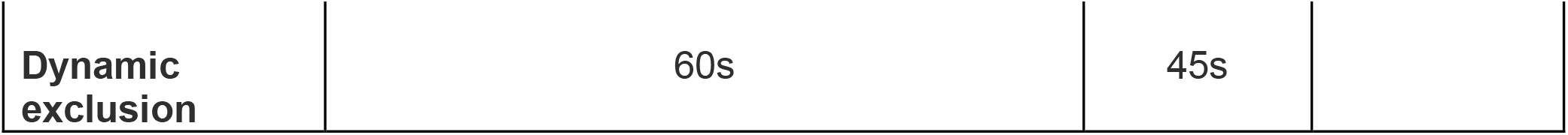
Basic LC and MS/MS parameter employed for data-dependent acquisition.

#### Adhesion analysis

Cells were harvested at 1000g for 30sec and subsequently resuspended in fresh TAP or HEPES/NMG buffer consisting of 5 mM HEPES, 1 mM KCl, 1 mM HCl, 200 µM EGTA and pH adjusted to 7.4 with N-methyl-D-glucamine. Reconstituted cells were following transferred to ibidi polymere channel slides with a height of 0.4mm. To give cells time to adhere and recover from centrifugation, mounted channels were incubated for 10 min before connecting them to the ibidi pump system and transfer to the microscopy chamber. Unidirectional flow was applied as soon as recording was started with 0.8 frames/sec using the 488nm laser at 0.5% power as light source and a 40x magnification. Cells measured in TAP medium were analysed at a constant flow rate of 3.77ml/min. Cells measured in Hepes/NMG devoid of calcium were subjected to computer controlled stepwise increasing pump pressure every 30 seconds (see table 4). Three biological replicates were performed. Error bars present standard deviation. Obtained videos were analysed using Fiji ImageJ^31^. Frames were transformed into binary images and normalised intensity plotted over time. Student t-test was performed at T50sec, T100sec, T150sec and T200sec.

**Table 4:** Computer controlled stepwise increasing pump pressure and corresponding calculated flowrate using ibidi pump system software.

| Pump pressure<br>[mbar] | Flowrate<br>[ml/min] | Duration<br>[sec] |
| --- | --- | --- |
| 5 | 3.77 | 25 |
| 10 | 7.38 | 30 |
| 20 | 14.18 | 30 |
| 40 | 26.28 | 30 |
| 60 | 36.72 | 30 |
| 80 | 45.83 | 30 |
| 95 | 51.94 | ∞ |

#### TIRF microscopy

TIRF microscopy was applied to assess gliding behaviour of *C. reinhardtii* strains expressing fluorescently tagged IFT46. Therefore, cell densities were adjusted to 10^5 cells·mL^−1^ in TAP. Samples were loaded to a glass bottom microscopy chamber and refreshed every 20 min while imaging. TIRF microscopy was performed at room temperature with a Nikon Eclipse Ti and a 100x objective. IFT46::YFP/mNG was excited at 488 nm and fluorescence was recorded with an iXon Ultra EMCCD camera (Andor). For analysis, images were captured with NIS-Elements software over 30 s at 10 fps.

#### Spinning-disk microscopy

Cells were deposited on a cover slide in a custom holder and imaged on an Olympus IX83 platform using an Olympus 100x/1.4 U Plan SApo Oil immersion objective and a Yokogawa CSU-W1 spinning disk scan head with two attached Ador iXon Ultra 888 cameras resulting in images with a pixel size of 79nm. Z-stacks were acquired using a piezo driven Prior Pro Scan III stage.

#### Microsphere binding assay

Equal number of cells of CC-124 *fmg1a-fmg1b* and FMG1A-mScarlet-FMG1B-mNeonGreen were merged and washed twice in TAP medium before an excess amount of thoroughly TAP washed polystyrene microspheres (0.2µm) were added to the cell suspension. After 15 min of incubation under gentle agitation in light, samples were fixed with 0.2% glutaraldehyde and directly subjected to microscopic analysis.

#### Single Particle Analysis

##### Protein Purification

FMG1-B was purified by affinity chromatography using a FPLC system (Cytiva Aktapure). Specifically, cilia were isolated from 20L of *fmg1-a* bubbling culture in 14h/10h light/dark to ensure the presence of only one isoform. Cells were concentrated for 10 min at 1200 RCF in at F8S-6×1000y rotor and resuspended in roughly 50 mL of 10 mM HEPES per liter of culture, then pooled and centrifuged again for 10 min at 1200 RCF before resuspending to a total of 1.5L with 10 mM HEPES 300uM CaCl_2_. Deciliation was induced by adding 11.25mL of 1M acetic acid while stirring, followed by neutralization with 11.25mL of 1M NaOH, 150mL of 50% sucrose as osmolyte, 6mL of 1M MgSO4 and 1.5mL of 500mM EDTA and 10mL of 100mM PMSF. Cells were then spun out in two centrifugation steps of 10 min at 1400 RCF and the supernatant containing the cilia was collected. Isolated cilia were then concentrated by centrifugation at 30 kRCF for 30 min at 4C and resuspended in ConA binding buffer (10mM HEPES pH 7.2, 500mM NaCl, 1mM MnCl2, 1mM CaCl2) to a total of 8mL, followed by membrane lysis with 0.1% final concentration IGEPAL CA-630. The lysate was clarified by 10min of centrifugation at 16 kRCF in 2mL tubes, followed by passing through a 0.22uM syringe filter. The clarified lysate was loaded onto a 1mL ConA sepharose 4B (Cytiva) column at 0.2mL/min, washed with 20mL of binding buffer + 0.1% IGEPAL followed by elution with a gradient of binding buffer + 0-1M sucrose over 20mL at 1mL/min. The purified FMG1-B was then concentrated to 1mL at ∼2 mg/mL. To polish and passivate the protein, a Superose 6 column equilibrated in imaging buffer (10mM HEPES, 300mM NaCl) was loaded with 1mL of imaging buffer + 10mg of Amphipol A8-35, followed directly by 500uL of FMG1-B. Peak fractions were collected and concentrated to 500uL at 0.8 mg/mL, then immediately plunge frozen on Quantifoil 1.2/1.3 grids at 1x, 2x and 4x dilution on a Leica GP2.

##### Data Acquisition and processing

4x diluted samples were screened on a Glacios and then imaged on a Krios G4 equipped with a Falcon4 camera. A total of 8010 images were acquired at 0.955 Å/px with a total dose of 70e/Å^2^. Single particle processing was performed in CryoSPARC^32^ unless otherwise noted. Raw eer movies were motion corrected, CTF estimated by patch CTF and particles were picked first by blob picking, classified and used to train crYOLO^33^ for particle picking. Picked positions were then extracted again, classified over multiple rounds with solvent clamping and per-particle scaling and an initial model was estimated de-novo. Non-uniform refinement of 3.6M particles yielded a density with 3.2A global FSC resolution, but due to the high flexibility of the structure, the local resolution of certain parts was much lower. 3Dflex refinement was performed with a manual segmentation using 9 latent dimensions and resulted in a more uniform refinement. This volume was used as reference for piecewise local refinement in 7 sub-volumes which were then combined to a final map with FSC resolution between 2.8-3.4A.

##### Model building

An initial guess for the peptide model was generated using AlphaFold3. The result was locally well predicted, but had an incorrect macro-structure. The initial prediction was manually manipulated to closer resemble the cryoEM density map in ChimeraX^34^, then relaxed under the density by coarse grain molecular dynamics using IModFit^10^. Finally, the model was fine-tuned on a per-residue basis by GPU assisted real time molecular dynamics in ChimeraX using ISOLDE^11^.

#### Cryo Electron Tomography

##### Data acquisition

Chlamydomonas cells were frozen directly on grid by applying a cell suspension and blotting away the excess before plunging into liquid ethane on a Leica GP2 plunge freezer. Grids were screened for cilia in open holes and acquired dose-symmetrically from −60 to 60 degrees in 2 degree increments. Images were acquired using using a Krios G4 with a Falcon 4 camera at 3.06Å/pix or 3.78Å/pix, 2.8 s exposure at a dose rate of 1.018e^−^/Å^2^/sec for a total of 2.85e^−^/Å^2^ per image.

##### Reconstruction

Pre-processing and tomogram reconstruction were done with the help of TOMOMAN. Raw EER files were motion-corrected in Relion, followed by CTF estimation with CTFFIND4 and finally automatic reconstruction at 4x binning with IMOD. For visualisation purposes tomograms were filtered with IsoNet^35^ deconvolution with a deconvolution strength and snrfalloff of 1. For tomogram segmentation (Fig. 1C), the glycocalyx was masked by fitting the membrane along the cilium with a custom python software with a closed spline which was then offset by a defined amount to get a mask containing the glycocalyx. Using this mask, densities and orientations matching a downscaled single-particle map of FMG1B were identified by template matching using GAPSTOP^36^ with GPU acceleration. Resulting peaks were filtered by filling in the predicted densities in a simulated tomogram while excluding particles with lower cross-correlation that would result in significant overlap. Resulting FMG1 positions were inspected in ChimeraX/ArtiaX^37^ and finally imported into Blender 4.1 where microtubules and membranes were manually modelled based on tomography slices.

### Quantification and statistical analysis

#### Gliding velocity and distance

Obtained TIRF videos were evaluated by use of Fiji ImageJ KymograpBuilder to determine gliding velocities and distances as described in^38^. All gliding events (velocity > 0µm/sec) were included. Gliding velocities of 40 cells (161 gliding events) in *ift46::NIT IFT46-mNG* and 42 cells (219 gliding events) in *fmg1b-fmg1a ift46::NIT IFT46-mNG* were obtained and merged from three biological replicates. In total 60 gliding distances (20 per strain and replicate) were evaluated using ImageJ^31^ measure. Man-Whitney-U test was performed for statistical evaluation.

#### Mass Spectrometry

For label-free protein quantification, if not further stated samples were measured in biological quadruplicates using standard, data dependent acquisition. Protein identification and quantification was performed using Fragpipe default LFQ settings excluding MBR and allowing unique and razor peptides for quantification. LFQ data (combined_proteins.tsv) was analysed using fragpipe-analyst developers version^39^ using MaxLFQ Intensity. Variance stabilising normalisation was allowed. Imputation was performed using perseus imputation and False discovery rate (FDR) correction was carried out using the Benjamini-Hochberg method. Quantification based on single peptide identifications as well as proteins identified in less than 70% of replicates per group were excluded. *N*-glycopeptide identification and validation from ISF-MS/MS runs was performed using Ursgal as described elsewhere^40^. Disulfide peptide identification and validation was performed using MaxQuantv2.0.3.1. N-terminal acetylation and methionine oxidation as well as carbamidomethylation of cysteines were allowed as variable modifications. Cysteine-Cysteine (−2.016 Da) was implemented as a potential cross-link.

## Conflict of interest

The authors declare that they have no conflict of interest.

**Data S1:**
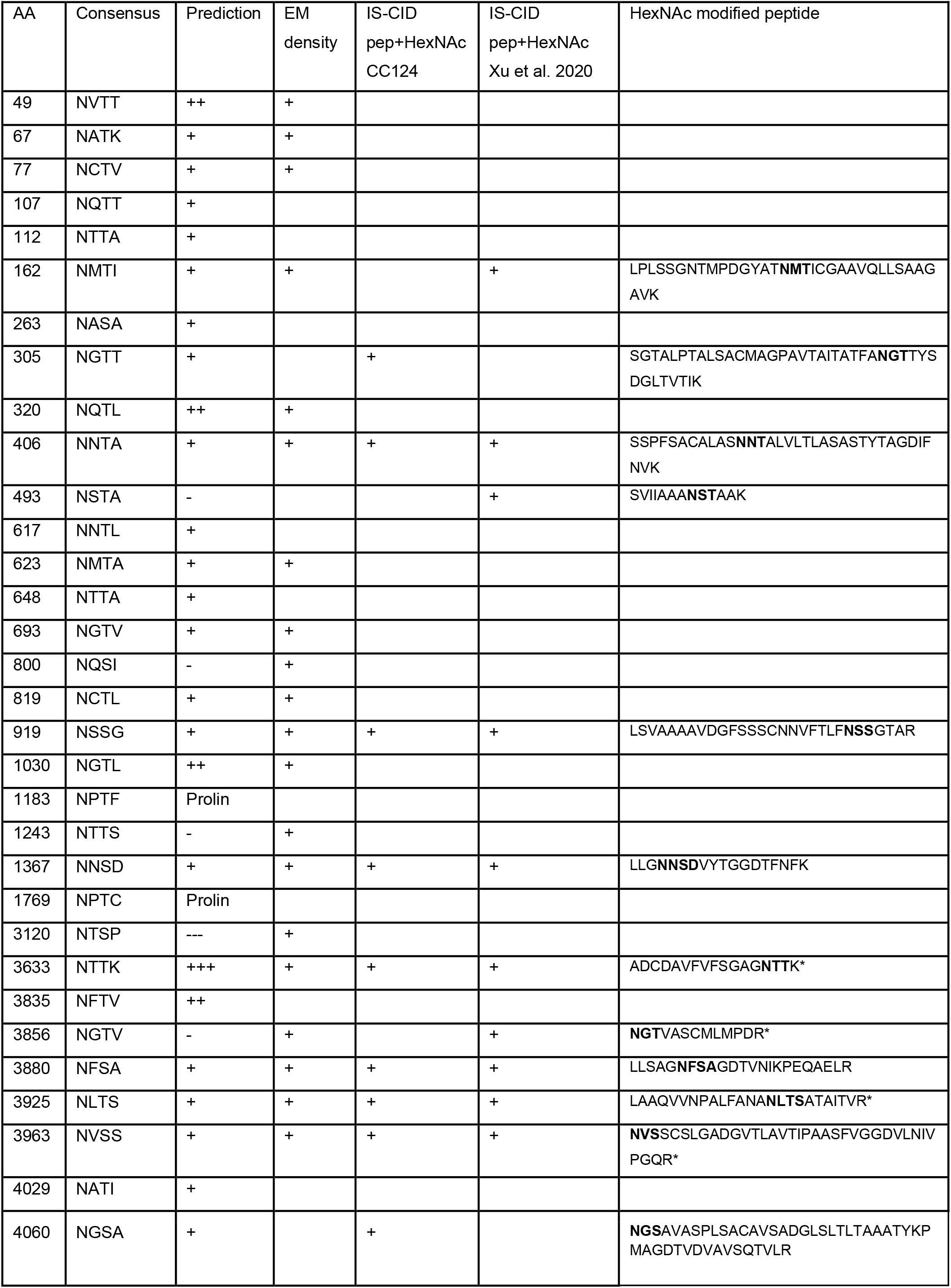
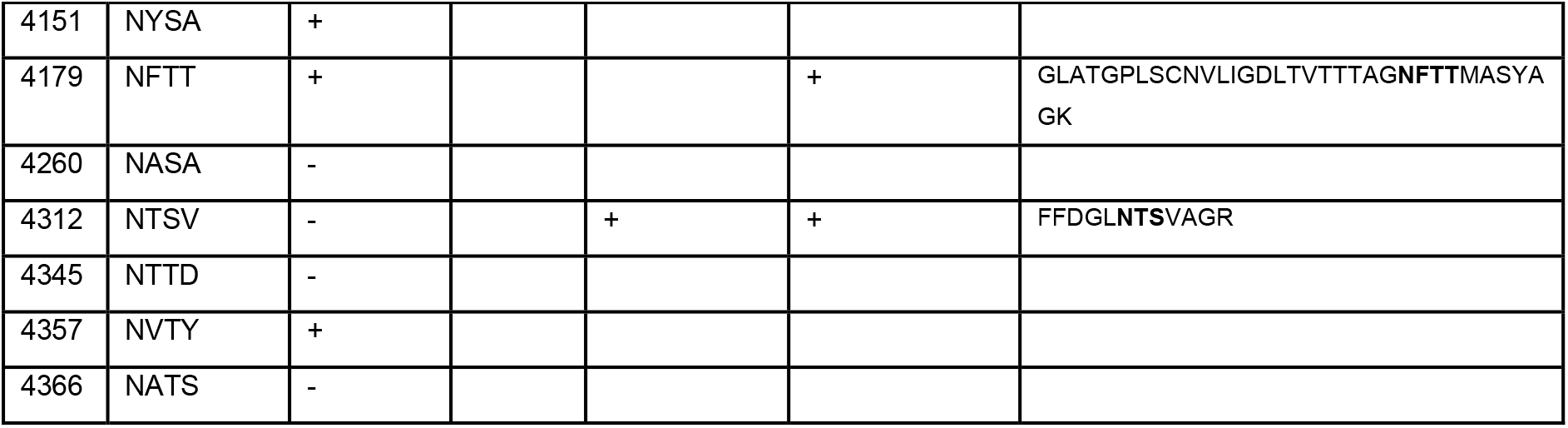
List of N-glycosites of FMG1B identified by computational and experimental methods. Net*N*-glyc predicted *N*-glycosites in FMG1B correlate greatly with additional densities observed in CryoEM, IS-CID-MS/MS identified glycopeptides in CC124 and identified glycopeptides for FMG1B in ^2^. Peptides marked with a * are indistinguishable from FMG1A. Related to Figure 3.

## Supplementary Video Legends

**Video S1: Flexible single particle refinement of FMG1B with 3DFlex**. Individual different latent dimensions are shown sequentially. Related to figure 3.

**Data S1:** Archive file containing code and genetic design information.

## References

1. Shih, S.M., Engel, B.D., Kocabas, F., Bilyard, T., Gennerich, A., Marshall, W.F., and Yildiz, A. (2013). Intraflagellar transport drives flagellar surface motility. eLife 2. 10.7554/elife.00744.

2. Xu, N., Oltmanns, A., Zhao, L., Girot, A., Karimi, M., Hoepfner, L., Kelterborn, S., Scholz, M., Beißel, J., Hegemann, P., et al. (2020). Altered N-glycan composition impacts flagella-mediated adhesion in Chlamydomonas reinhardtii. eLife 9. 10.7554/elife.58805.

3. Möckl, L. (2020). The Emerging Role of the Mammalian Glycocalyx in Functional Membrane Organization and Immune System Regulation. Front. Cell Dev. Biol. 8, 253. 10.3389/fcell.2020.00253.

4. Bloodgood, R.A., Tetreault, J., and Sloboda, R.D. (2019). The Chlamydomonas flagellar membrane glycoprotein FMG-1B is necessary for expression of force at the flagellar surface. J. Cell Sci. 132. 10.1242/jcs.233429.

5. Pazour, G.J., Agrin, N., Leszyk, J., and Witman, G.B. (2005). Proteomic analysis of a eukaryotic cilium. J. Cell Biol. 170, 103–113. 10.1083/jcb.200504008.

6. Li, X., Patena, W., Fauser, F., Jinkerson, R.E., Saroussi, S., Meyer, M.T., Ivanova, N., Robertson, J.M., Yue, R., Zhang, R., et al. (2019). A genome-wide algal mutant library and functional screen identifies genes required for eukaryotic photosynthesis. Nat. Genet. 51, 627–635. 10.1038/s41588-019-0370-6.

7. Bloodgood, R.A., Salomonsky, N.L., and Reinhart, F.D. (1987). Use of carbohydrate probes in conjunction with fluorescence-activated cell sorting to select mutant cell lines of Chlamydomonas with defects in cell surface glycoproteins. Exp. Cell Res. 173, 572–585. 10.1016/0014-4827(87)90296-5.

8. Mathieu-Rivet, E., Scholz, M., Arias, C., Dardelle, F., Schulze, S., Le Mauff, F., Teo, G., Hochmal, A.K., Blanco-Rivero, A., Loutelier-Bourhis, C., et al. (2013). Exploring the N-glycosylation Pathway in Chlamydomonas reinhardtii Unravels Novel Complex Structures. Mol. Cell. Proteomics 12, 3160–3183. 10.1074/mcp.m113.028191.

9. Abramson, J., Adler, J., Dunger, J., Evans, R., Green, T., Pritzel, A., Ronneberger, O., Willmore, L., Ballard, A.J., Bambrick, J., et al. (2024). Accurate structure prediction of biomolecular interactions with AlphaFold 3. Nature 630, 493–500. 10.1038/s41586-024-07487-w.

10. Lopéz-Blanco, J.R., and Chacón, P. (2013). iMODFIT: Efficient and robust flexible fitting based on vibrational analysis in internal coordinates. J. Struct. Biol. 184, 261–270. 10.1016/j.jsb.2013.08.010.

11. Croll, T.I. (2018). ISOLDE: a physically realistic environment for model building into low-resolution electron-density maps. Acta Crystallogr. Sect. Struct. Biol. 74, 519–530. 10.1107/s2059798318002425.

12. Gupta, R., and Brunak, S. (2001). Prediction of glycosylation across the human proteome and the correlation to protein function. In Biocomputing 2002 (WORLD SCIENTIFIC). 10.1142/9789812799623_0029.

13. Craig, R.J., Hasan, A.R., Ness, R.W., and Keightley, P.D. (2021). Comparative genomics of Chlamydomonas. Plant Cell 33, 1016–1041. 10.1093/plcell/koab026.

14. Pajic, P., Shen, S., Qu, J., May, A.J., Knox, S., Ruhl, S., and Gokcumen, O. (2022). A mechanism of gene evolution generating mucin function. Sci. Adv. 8. 10.1126/sciadv.abm8757.

15. Pino, V., Ramsauer, V.P., Salas, P., Carraway, C.A.C., and Carraway, K.L. (2006). Membrane Mucin Muc4 Induces Density-dependent Changes in ERK Activation in Mammary Epithelial and Tumor Cells. J. Biol. Chem. 281, 29411–29420. 10.1074/jbc.m604858200.

16. Proksch, J., Dal Colle, M.C.S., Heinz, F., Schmidt, R.F., Gottwald, J., Delbianco, M., Keller, B.G., Gradzielski, M., Alexiev, U., and Koksch, B. (2024). Impact of glycan nature on structure and viscoelastic properties of glycopeptide hydrogels. J. Pept. Sci. 30. 10.1002/psc.3599.

17. Dai, J., Ma, M., Niu, Q., Eisert, R.J., Wang, X., Das, P., Lechtreck, K.F., Dutcher, S.K., Zhang, R., and Brown, A. (2024). Mastigoneme structure reveals insights into the O-linked glycosylation code of native hydroxyproline-rich helices. Cell 187, 1907–1921.e16. 10.1016/j.cell.2024.03.005.

18. Huang, J., Tao, H., Chen, J., Shen, Y., Lei, J., Pan, J., Yan, C., and Yan, N. (2024). Structure-guided discovery of protein and glycan components in native mastigonemes. Cell 187, 1733–1744.e12. 10.1016/j.cell.2024.02.037.

19. The Chlamydomonas ciliary membrane and its dynamic properties (2023). In The Chlamydomonas Sourcebook (Elsevier), pp. 247–271. 10.1016/b978-0-12-822508-0.00005-8.

20. Betleja, E. (2012). Analysis of the gliding machinery in the green alga, Chlamydomonas reinhardtii.

21. Kreis, C.T., Le Blay, M., Linne, C., Makowski, M.M., and Bäumchen, O. (2018). Adhesion of Chlamydomonas microalgae to surfaces is switchable by light. Nat. Phys. 14, 45–49. 10.1038/nphys4258.

22. Kamiya, R., Shiba, K., Inaba, K., and Kato-Minoura, T. (2018). Release of Sticky Glycoproteins from Chlamydomonas Flagella During Microsphere Translocation on the Surface Membrane. Zoolog. Sci. 35, 299. 10.2108/zs180025.

23. Das, P., Mekonnen, B., Alkhofash, R., Ingle, A., Workman, E.B., Feather, A., Liu, P., and Lechtreck, K.F. (2023). Small Interactor of PKD2 (SIP), a novel PKD2-related single-pass transmembrane protein, is required for proteolytic processing and ciliary import ofChlamydomonasPKD2. Preprint at Cold Spring Harbor Laboratory, 10.1101/2023.06.13.544839 10.1101/2023.06.13.544839.

24. Seifert, G.J. (2018). Fascinating Fasciclins: A Surprisingly Widespread Family of Proteins that Mediate Interactions between the Cell Exterior and the Cell Surface. Int. J. Mol. Sci. 19, 1628. 10.3390/ijms19061628.

25. Willis, A., Eason-Hubbard, M., Hodson, O., Maheswari, U., Bowler, C., and Wetherbee, R. (2014). Adhesion molecules from the diatom Phaeodactylum tricornutum (Bacillariophyceae): genomic identification by amino-acid profiling and in vivo analysis. J. Phycol. 50, 837–849. 10.1111/jpy.12214.

26. Sasso, S., Stibor, H., Mittag, M., and Grossman, A.R. (2018). From molecular manipulation of domesticated Chlamydomonas reinhardtii to survival in nature. eLife 7. 10.7554/elife.39233.

27. Nievergelt, A.P., Diener, D.R., Bogdanova, A., Brown, T., and Pigino, G. (2024). Protocol for precision editing of endogenous Chlamydomonas reinhardtii genes with CRISPR-Cas. STAR Protoc. 5, 102774. 10.1016/j.xpro.2023.102774.

28. Nievergelt, A.P., Diener, D.R., Bogdanova, A., Brown, T., and Pigino, G. (2023). Efficient precision editing of endogenous Chlamydomonas reinhardtii genes with CRISPR-Cas. Cell Rep. Methods 3, 100562. 10.1016/j.crmeth.2023.100562.

29. Witman, G.B., Carlson, K., Berliner, J., and Rosenbaum, L. (1972). CHLAMYDOMONAS FLAGELLA I. Isolation and Electrophoretic Analysis ofMicrotubules, Matrix, Membranes, and Mastigonemes. ThE JOURNAL OF CELL BIOLOG.

30. Rappsilber, J., Mann, M., and Ishihama, Y. (2007). Protocol for micro-purification, enrichment, pre-fractionation and storage of peptides for proteomics using StageTips. Nat. Protoc. 2, 1896–1906. 10.1038/nprot.2007.261.

31. Schindelin, J., Arganda-Carreras, I., Frise, E., Kaynig, V., Longair, M., Pietzsch, T., Preibisch, S., Rueden, C., Saalfeld, S., Schmid, B., et al. (2012). Fiji: an open-source platform for biological-image analysis. Nat. Methods 9, 676–682. 10.1038/nmeth.2019.

32. Punjani, A., Rubinstein, J.L., Fleet, D.J., and Brubaker, M.A. (2017). cryoSPARC: algorithms for rapid unsupervised cryo-EM structure determination. Nat. Methods 14, 290–296. 10.1038/nmeth.4169.

33. Wagner, T., Merino, F., Stabrin, M., Moriya, T., Antoni, C., Apelbaum, A., Hagel, P., Sitsel, O., Raisch, T., Prumbaum, D., et al. (2019). SPHIRE-crYOLO is a fast and accurate fully automated particle picker for cryo-EM. Commun. Biol. 2, 218. 10.1038/s42003-019-0437-z.

34. Pettersen, E.F., Goddard, T.D., Huang, C.C., Meng, E.C., Couch, G.S., Croll, T.I., Morris, J.H., and Ferrin, T.E. (2021). UCSF ChimeraX : Structure visualization for researchers, educators, and developers. Protein Sci. 30, 70–82. 10.1002/pro.3943.

35. Liu, Y.-T., Zhang, H., Wang, H., Tao, C.-L., Bi, G.-Q., and Zhou, Z.H. (2022). Isotropic reconstruction for electron tomography with deep learning. Nat. Commun. 13, 6482. 10.1038/s41467-022-33957-8.

36. Cruz-León, S., Majtner, T., Hoffmann, P.C., Kreysing, J.P., Kehl, S., Tuijtel, M.W., Schaefer, S.L., Geißler, K., Beck, M., Turoňová, B., et al. (2024). High-confidence 3D template matching for cryo-electron tomography. Nat. Commun. 15, 3992. 10.1038/s41467-024-47839-8.

37. Ermel, U.H., Arghittu, S.M., and Frangakis, A.S. (2022). ArtiaX : An electron tomography toolbox for the interactive handling of s ub-tomograms in UCSF ChimeraX. Protein Sci. 31, e4472. 10.1002/pro.4472.

38. In vivo Imaging of IFT in Chlamydomonas Flagella (2013). In Methods in Enzymology (Elsevier), pp. 265–284. 10.1016/b978-0-12-397945-2.00015-9.

39. Shah, A.D., Goode, R.J.A., Huang, C., Powell, D.R., and Schittenhelm, R.B. (2020). LFQ-Analyst: An Easy-To-Use Interactive Web Platform To Analyze and Visualize Label-Free Proteomics Data Preprocessed with MaxQuant. J. Proteome Res. 19, 204–211. 10.1021/acs.jproteome.9b00496.

40. Schulze, S., Oltmanns, A., Fufezan, C., Krägenbring, J., Mormann, M., Pohlschröder, M., and Hippler, M. (2021). SugarPy facilitates the universal, discovery-driven analysis of intact glycopeptides. Bioinformatics 36, 5330–5336. 10.1093/bioinformatics/btaa1042.

